# Recruitment of the protein phosphatase-1 catalytic subunit to promoters by the dual-function transcription factor RFX1

**DOI:** 10.1101/493429

**Authors:** Yoav Lubelsky, Yosef Shaul

## Abstract

RFX proteins are a family of conserved DNA binding proteins involved in various, essential cellular and developmental processes. RFX1 is a ubiquitously expressed, dual-activity transcription factor capable of both activation and repression of target genes.

The exact mechanism by which RFX1 regulates its target is not known yet. In this work, we show that the C-terminal repression domain of RFX1 interacts with the Serine/Threonine protein phosphatase PP1c, and that interaction with RFX1 can target PP1c to specific sites in the genome. Given that PP1c was shown to de-phosphorylate several transcription factors, as well as the regulatory C-terminal domain of RNA Polymerase II the recruitment of PP1c to promoters may be a mechanism by which RFX1 regulates the target genes.

## Introduction

Protein phosphorylation plays a major role in the regulation of numerous cellular functions. The specific phosphorylation of various target proteins is mediated by a large number of protein kinases each with a different specificity [1]. The phosphorylated sequence can then be used as a docking site for other proteins, lead to changes in the activity or sub-cellular location of the phosphorylated protein, or influence the protein stability by targeting it to or protecting it from proteasomal degradation [2-6].

In contrast to the large number of protein kinases the number of protein phosphatases, whose role is to de-phosphorylate proteins, is very small and their sequence specificity is very broad [7-9]. Protein serine/threonine phosphatases are classified into three families PPM, FCP and PPP. The PPP family, which is the most abundant, is further divided into PP1, PP2A, PP2B, and PP5 subfamilies [10, 11]. The PP1 family is constituted of 4 isoforms of the PP1 catalytic subunit (PP1c) PP1cα, PP1cβ/δ, PP1cγ1 and PP1cγ2 that are expressed from 3 genes. With the exception of the testis specific PP1cγ2 all isoforms are ubiquitously expressed [12].

By itself the PP1 catalytic subunit is promiscuous and substrate specificity is achieved by a large number of interacting proteins that serve as regulatory subunits, regulating its activity and/or targeting PP1c to different substrates or subcellular locations [13-16]. While the different PP1c interacting-proteins do not share any overall sequence similarity most of them contain the consensus PP1c interacting motif R/K-X_0-1_-V/I-[P]-F/W motif, commonly referred to as the RVxF motif, where X is any amino acid and [P] is any amino acid except proline, which interact with the hydrophobic channel of the catalytic subunit [17, 18].

RFX1 is a member of the evolutionally conserved family of DNA binding proteins, characterized by a unique winged helix DNA binding domain, which is highly conserved in eukaryotic organisms [19-21]. The known RFX family includes one member in *Sacchromyces cerevisiae, Schizosacchromyces pombe, Caenorhabditis elegans* and the fungus *Acremonium chrysogenum*, two members in *Drosophila melanogaster*, and eight members in mammals [20, 22-27].

RFX1 has a transcription repression and a transcription activation domain that can neutralize each other in exogenous systems [28]. RFX1 was originally identified as an activator of Hepatitis B gene expression [29]. On the other hand, RFX1 was shown to represses the expression of genes such as PCNA [30], c-myc [31], the RNR R2 subunit and its own gene [32]. It has been suggested that the function of RFX1 in supporting or repressing transcription depends on the promoter context [28].

In this work, we show that PP1c is recruited to promoters by RFX1. PP1c de-phosphorylates and subsequently affect the function of transcription factor as well as components of the general transcription machinery [33-39]. We suggest a model for the context dependent function of RFX1 which is achieved by recruiting PP1c to promoters leading to the de-phosphorylation of neighboring factors.

## Materials and methods

### Cell culture and transfections

All cells were grown in Dulbecco’s Modified Eagle Medium (DMEM) (Gibco) containing 100U/ml penicillin and 100μg/ml Streptomycin, supplemented with 9% fetal calf serum. Transfections were done by the calcium phosphate precipitation method or using Polyethyleneimine (PEI linear, M*r* 25000 from Polyscience Inc.) [40].

### Reporter gene assay

RFX1-luciferase reporters, Gal4-DBD fusion, and HA-RFX1 constructs were previously described [28, 32]. In order to generate the RVxFm construct R780 and F783 were changed to alanine using site directed mutagenesis. Luciferase expression was normalized to Ef1a-renilla expression. GFP-PP1 plasmids [41] were a gift from Mathieu Bollen.

### Chromatin Immunoprecipitation

Chromatin immunoprecipitation (ChIP) was performed according to the protocol of Ainbinder et. al. [42]. Briefly, formaldehyde crosslinked protein-DNA complexes were precipitated by incubation overnight with anti-RFX1 polyclonal antibody or with rabbit pre-immune serum as a negative control. The extract was then cleared by centrifugation and incubated for an additional 3 hours with protein-A conjugated sepharose beads (Pharmacia). Precipitated DNA fragments were extracted and amplified with specific primers. For ChIP of target promoters located on a plasmid the protocol was slightly modified. The number of sonication pulses was reduced and the extract was diluted 5 times instead of 10 times in the dilution buffer. Precipitated chromatin was detected by end point PCR or by qPCR using the Lightcycler-480 (Roche).

## Results

### RFX1 mediate repression by recruiting an HDAC independent co-repressor

In yeast the RFX homologue Crt1p represses the expression of its targets by recruiting the Ssn6/Tup1 general repressor complex [43]. The Ssn6/complex as well as its mammalian counterpart the TLE protein family represses transcription by recruiting Histone de-acetylase proteins [44-46]. The repression domain of RFX1 is located in the C-terminus end of the protein, while the N-terminus contains the activation domain and these domains are capable of neutralizing each other [28]. In our previous work [32], we have shown that RFX1 represses its own promoter possibly by recruiting of an unknown endogenous co-repressor.

To investigate the nature of the corepressor we overexpressed fragments of RFX1 to serve as a decoy for the presumed corepressor. The cells were transfected with a luciferase reporter under the control of the RFX1 promoter [32] in the presence of different fragments of RFX1 fused to the Gal4 DNA binding domain [28] (fig 1A). Only the entire C-terminus (aa 529-979) fragment significantly increased RFX1 promoter activity (fig 1B) indicating that the whole C-terminal fragment is involved in the recruitment of the co-repressor. Overexpressing the wtRFX1 (Fig 1B, white bars) was able to restore repression further supporting the competition model In order to test the involvement of histone de-acetylases corepressors in RFX1 mediated repression we treated cells transfected with the RFX1 promoter luciferase reporter [32] with either Trichostatin A (TSA), an inhibitor of class I and II HDACs or with Nicotinamide (NA), an inhibitor of class III HDACs. The expression level of the reporter was elevated after treatment with both inhibitors indicating that histone acetylation effects the expression of the RFX1 promoter (fig 1C black bars).

**Figure 1:**
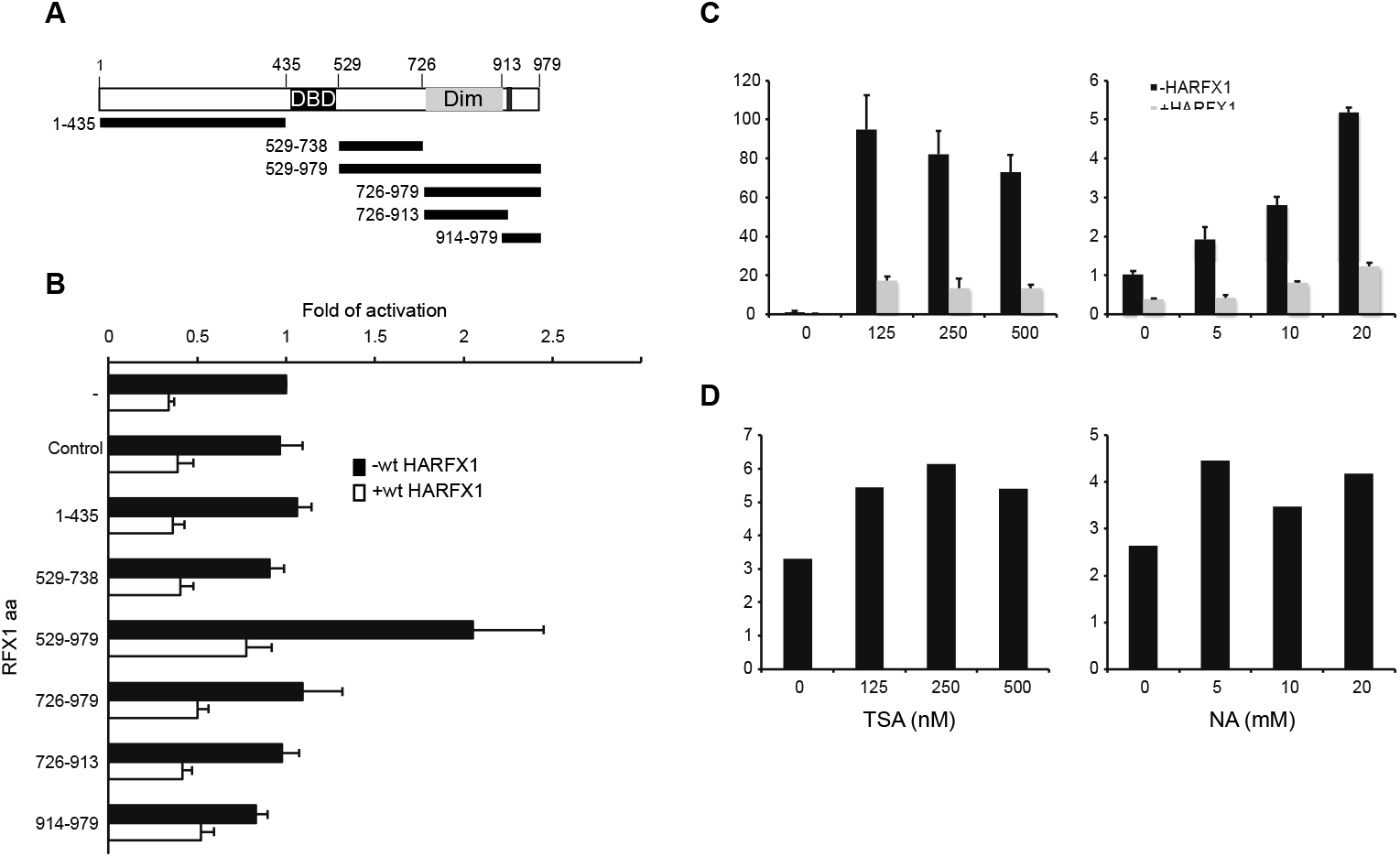
RFX1 transcription repression is mediated through its C-terminus. **A:** schematic drawing of HARFX1 with the DNA-binding domain (DBD) and dimerization domain (Dim). The black bars indicate the different RFX1 regions expressed in B. **B:** Activity of the RFX1-luc reporter in the presence (+) or absence (-) of co-transfected HARFX1 and the different Gal4-RFX1 constructs. The number of the amino acids of RFX1 fragment in each of the Gal4-chimeric construct are indicated. **C:** Luciferase activity from cells transfected with the RFX1-luc reporter with (grey bars) or without (black bars) HARFX1 and treated for 24 hours with the indicated concentration of either Trichostatin A (TSA) or Nicotine amide (NA). Y-axis show the fold of induction over untreated cells **D:** The level of repression by RFX1 in panel C is calculated as the ratio between the activity of the reporter in the absence or presence of RFX1 overexpression.

However, over expression of RFX1 repressed the reporter expression in the presence of the inhibitors (fig 1C grey bars) and the level of the repression was not affected by inhibitors concentration (fig 1D). This data suggests that RFX1-mediated repression is achieved by some other mechanism that is dominant over HDAC activity.

### RFX1 binds Protein Phosphatase 1

Previous work in our lab has identified a phosphatase activity that is co-purified with RFX1, the RFX1 associated phosphatase was identified as PP1 (Hagar Greif, PhD thesis [47]). The PP1 catalytic subunits are promiscuous serine/threonine protein phosphatases that are targeted to specific substrate and/or cellular locations by a large number of interacting proteins [13, 14]. While the different PP1 interacting-proteins share no overall sequence similarity, most of them contain the consensus PP1 binding R/K-Xo_-1_-V/I-[P]-F/W motif, commonly referred to as the RVxF motif, where X is any amino acid and [P] is any amino acid except proline [17]. Several publications in recent years have shown interaction of PP1 with various regulators of transcription such as pTEF-b, and RNApol II [38, 48], as well as transcription factors [33, 37, 41, 49], raising the intriguing possibility that RFX1 might regulate the transcription of its target genes by recruiting PP1c.

Inspection of the sequence revealed that RFX1 contains an RVxF motif between residues 780 and 783 that is conserved in other mammalian RFX1 proteins in addition a second, albeit less characterized F-x-x-R/K-x-R/K motif [50] is located at aa 534-539 (Fig 2A) which may explain why the entire C-terminus is needed for efficient competition with the endogenous RFX1 (fig 1B). Both motifs are conserved throughout vertebrates (fig 1A, Supplementary figS1). To determine whether RFX1 can bind to PP1, we transfected HEK293 cells with GFP-tagged isoforms of the PP1 catalytic subunit [41, 51] and performed immunoprecipitation using RFX1 specific antibodies (Fig 2B). The RFX1 antibodies were able to precipitate GFP-PP1α and GFP-PP1γ1 (Fig 2B lanes 5 and 7) but not GFP-PP1β (Fig 2C lane 6) or GFP without any fusion protein that was used as negative control (Fig 2B lane 8).

**Figure 2:**
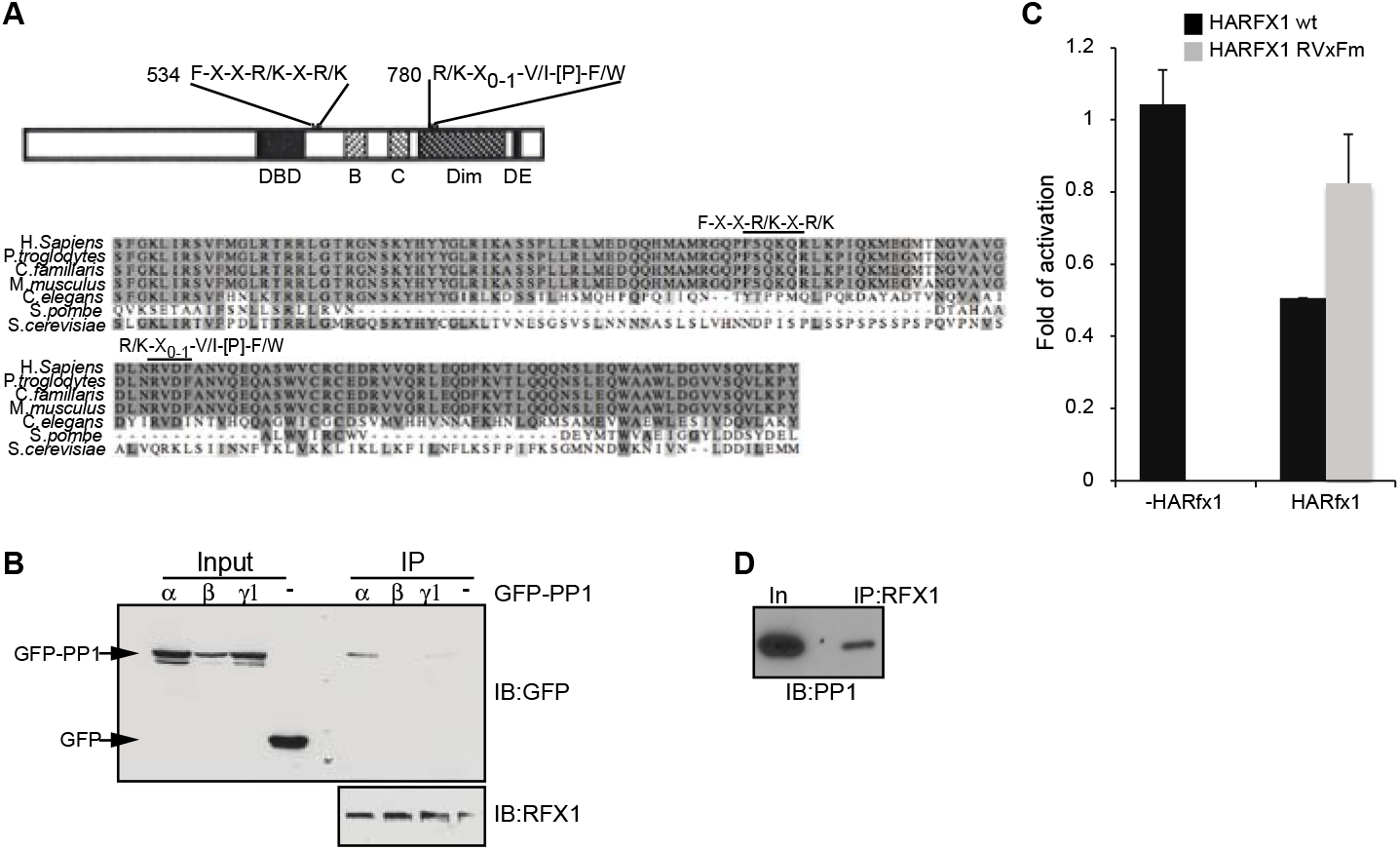
Rfx1 binds the PP1 catalytic subunit. **A:** Schematic draw of the RFX1 protein with the putative PP1 interaction motifs marked. Alignment of the region of RFX1 containing those motifs is shown below, the black line marks the motifs. **B:** PP1 interacts with RFX1. HEK293 cells were transfected with either pEGFP-PP1α, pEGFP-PP1β, pEGFP-PP1γ1 or pEGFP-C1 (empty GFP vector). Protein complexes were immunopercipitated using anti-RFX1 antibodies. Total extracts (Input) and Immunoprecipitated proteins (IP) were separated by SDS-PAGE and immunoblotted with anti-GFP (top) and anti-RFX1 (bottom) antibodies. **C:** Mutation in the RVxF PP1 interacting motif reduces RFX1 repressor activity. Activity of the RFX1-luc reporter in the absence of HARFX1 overexpression (left) or the presence HARFX1 wt (black) or HARFX1 RVxFm (grey). **D:** RFX1 interacts with the endogenous PP1. proteins from MCF7 cells whole cell extract were Immunoprecipitated with anti-RFX1 antibodies and blotted with anti-PP1c antibodies.

To verify that the interaction between RFX1 and PP1c occur at the level of the endogenous proteins as well, RFX1 was immunoprecipitated from MCF7 cells and probed with an antibody for the PP1 catalytic subunit. PP1c was precipitated with the anti-RFX1 antibodies (fig 2C), indicating that RFX1 interacts with endogenous PP1c. To examine the possible role of the PP1c interacting motif in RFX1 mediated repression we generated the an RFX1 mutant lacking the functional RVxF PP1c binding motif in which the conserved arginine (R780) and phenylalanine (F783) in the motif were changed to alanine. The ability of this mutant to repress the expression of the RFX1 promoter reporter was attenuated (fig 2D) supporting the involvement of PP1c in RFX1 mediated repression. These data suggest that repression by RFX1 is mediated by the recruitment of PP1c.

### PP1c is recruited to DNA by RFX1

Next we examined whether PP1c can be recruited to the DNA via its interaction with RFX1, for that we have performed ChIP assay in MCF7 cells with antibodies against RFX1, and PP1c as well as by using Microcystin fused agarose beads. The precipitated chromatin was tested for the RFX1 promoter, a known target of RFX1 [32]. RFX1 promoter was enriched in the RFX1 and PP1c immunoprecipitation and in the Mycrocystine pulldown (fig 3A black bars), no enrichment was detected in the unrelated GAPDH transcription start site (GAPDH TSS fig 3A, grey bars). Mycrocystine is a PP1c inhibitor which interacts with the active site of the enzyme [52]. The success in pull down of RFX1 target DNA by Microcystine beads indicates that the active site of RFX1-associated PP1c is not blocked and functionally persistent.

**Figure 3:**
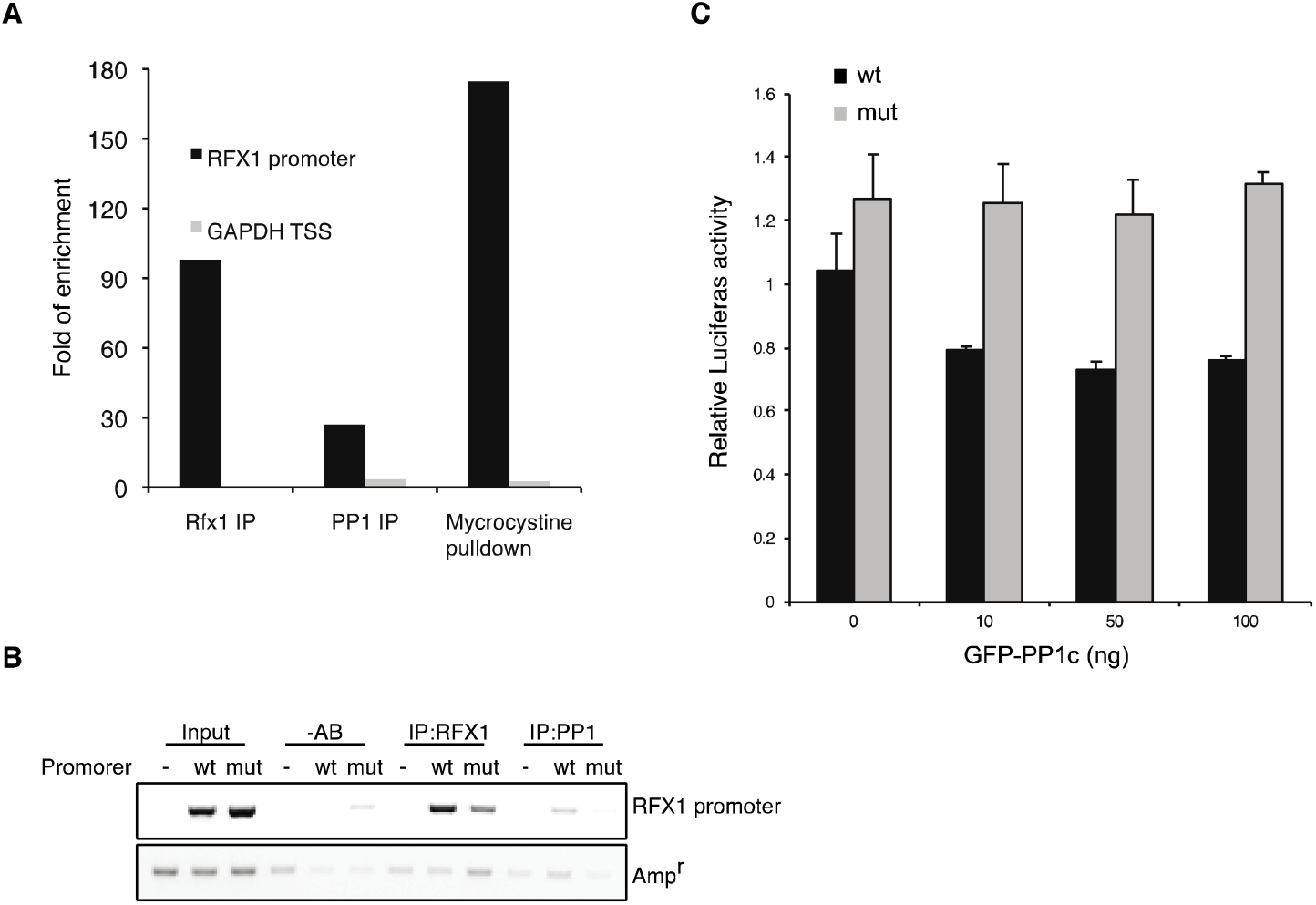
PP1c is associated with DNA via RFX1. **A: PP1 interacts with the RFX1 promoter**. Chromatin was precipitated from MCF7 cells using anti-RFX1 antibodies, anti-PP1c antibodies, and mycrocystine-agarose beads. The presence of the RFX1 promoter region and of a control region around the GAPDH transcription start site (GAPDH TSS) was analyzed by qPCR. The results are presented as fold of enrichment over the no antibody control IP. **B:** The interaction of PP1 with the RFX1 promoter is dependent on the RFX1 binding sites. MCF7 cells were transfected with plasmids containing no promoter, a wild type version of the *RFX1* promoter (wt) or a version of the *RFX1* promoter in which the RFX1 binding sites were mutated (mut). Chromatin was immunoprecipitated with either RFX1 or PP1 antibodies and the level of the RFX1 promoter or an unrelated part of the plasmid (the Ampicillin resistance gene) was analyzed by PCR. **C:** Repression by PP1c overexpression is dependent on RFX1. MCF7 cells were transfected with wt or mutant *RFX1* promoter reporter in the presence of increasing amount of GFP-PP1c. PP1c overexpression could repress the expression of wt reporter (black bars) but not of the mutant reporter (grey bars).

In order to verify that the association of PP1c with the DNA is indeed mediated by RFX1 we transfected MCF7 cells with reporter plasmid containing the RFX1 promoter (pGL3-RFX1 wt) or a plasmid containing the RFX1 promoter in which both RFX1 binding sites were mutated (pGL3-RFX1 mut) [32]. Formaldehyde crosslinked DNA was immunoprecipitated from these cells using RFX1 and PP1c antibodies, and analyzed for the cloned promoter (fwd primer from the cloned promoter region and rev primer from the luciferase ORF) or for an unrelated region of the vector (the Ampicilin resistance gene Amp^r^). Anti-RFX1 antibodies brought down the wild type promoter and to much lower level the mutant promoter due to the residual binding of RFX1 to the mutant sequence, as previously reported [32]. Interestingly, PP1 antibody pulled down the wt promoter, but neither the mutant promoter nor the empty vector (pGL3-Basic) (fig 3B), indicating that PP1c is recruited to the RFX1 promoter via RFX1.

Finally, we examined the RFX1 reporter in response to PP1c overexpression. The mutant reporter that only poorly bind RFX1 is more active than the wild-type possibly due to the endogenous RFX1 activity. Overexpression of GFP-PP1c further repressed the wt reporter but had no effect on the mutant (Fig 3C). These data suggest that RFX1 represses transcription of the target genes via the recruitment of the PP1c.

## Discussion

The RFX family of protein is highly conserved in evolution [20]. RFX1 was originally identified as an activator of HBV gene expression [29], however all the cellular target described to date are repressed by RFX1 [30-32, 53, 54]. We investigated the mode of autorepression of RFX1. Here we report that at least part of its repression activity is mediated by the recruited PP1c, the catalytic subunit of the protein phosphatase 1. This conclusion is based on the observations that RFX1 physically binds PP1c and that RFX1-PP1c complex can be detected at the RFX1 promoter region. Previously two RFX1 binding sites were identified at the RFX1 promoter [32]. Interestingly an RFX1 promoter mutant at these sites demonstrated both reduced RFX1-PP1c association and repression activity, suggesting that PP1c association and transcription repression are coupled.

The yeast RFX protein, Crt1p, represses transcription by recruiting the Ssn6/Tup1 complex [43], the homologue of the mammalian TLE proteins. We have shown that the signaling pathways and target genes are conserved between the Crt1 and RFX1 [32], however in this work we show that RFX1 represses transcription in an HDAC independent mechanism (fig 1C-D) that is mediated by the RFX1-PP1c complex. In agreement with this model the PP1c binding motifs in RFX1 are highly conserved in all vertebrate species but not in yeast (fig S1), the motifs are also missing from fly and worm RFX1 orthologues suggesting that they are likely to use a PP1 independent mechanism for repression, either an HDAC mechanism, similar to that described in yeast or a yet unidentified mechanism.

RFX1 is highly conserved and single nucleotide polymorphisms (SNPs) that are identified in >1% of samples (dbSNP 150, common SNPs) are restricted to regions outside the coding sequence (fig S2, top). With coding-sequence SNPs being rare it is difficult to assess their significance however SNPs that overlap the PP1c binding motifs do not change the amino-acid sequences while some adjacent SNPs that are still part of a highly conserved part of the proteins sometime do (fig S2, bottom). The exception is SNP #rs372349612 which on rare occasions (<0.001%) change F783 to L, importantly a peptide display screen for PP1c interacting sequences reveled that leucine can be tolerated in this position [55], adding further support for the importance of PP1c interaction for RFX1.

The serine/threonine phosphatase PP1 is a ubiquitously expressed enzyme. The PP1 catalytic subunit interacts with a large number of regulatory subunits that target it to specific locations and substrates (reviewed in [13, 14]). The involvement of PP1 in the regulation of transcription was previously described. PP1 can de-phosphorylate the C-terminal domain (CTD) of RNA polymerase II independently of FCP1, the canonical CTD phosphatase [38]. The regulated phosphorylation and de-phosphorylation of serine 2 and 5 in the CTD repeats is crucial for initiation of transcription and for the switch between initiation and elongation [56].

Searching the human proteome for occurrences of the PP1c binding RVxf motif identified possible sites in 4801 proteins in the Swissprot database, 134 of them were annotated as being involved in transcription (Supplementary table 1), gene ontology analysis revealed that the potential PP1c binding transcription regulators are enriched for Viral pathways including HBV, a known RFX1 target (Supplementary table 2). In fact, viruses exploit PP1 for regulating their genes as has been demonstrated for HIV [41].

Certain transcription factors such as p53 and CREB are regulated by PP1 but not involved directly in recruiting PP1, like RFX1. For example, PP1 de-phosphorylate p53 on serine 15 and 37 but the mechanism of PP1 recruitment and the function of de-phosphorylation is not yet clear [33]. Interestingly the presence of an RFX1 site next to the p53 binding site, as has been demonstrated in the context of the HBV enhancer, turns p53 from an activator to a repressor of transcription [57] raising the possibility that RFX1 change the function of p53 by recruiting PP1c to its vicinity and subsequently leading to its de-phosphorylation. Whether a similar mechanism regulates some other genes is an important question to be investigated.

## Supporting information

## Supplementary information

**Supplementary Figure S1:**
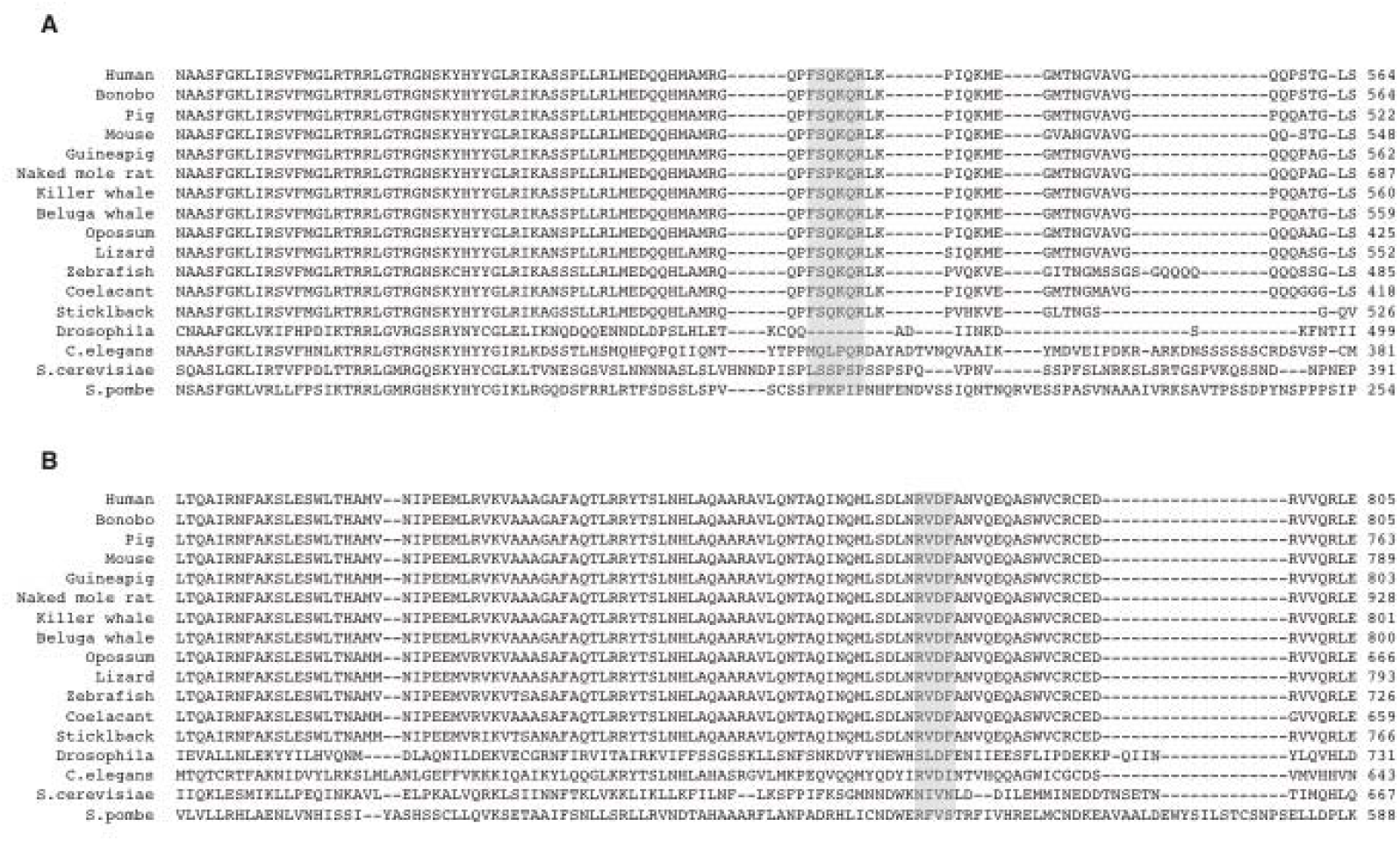
Multiple alignment of RFX1 sequences from different organisms. Alignment was preformed using the Clustal Omega web tool [1] (https://www.ebi.ac.uk/Tools/msa/clustalo/) Shaded regions indicate the FXXRXR (A) and RVXF (B) PP1binding motifs. Protein sequences used and the full alignment are in Supplementary data file (SuppData1.pdf).

**Supplementary Figure S2:**
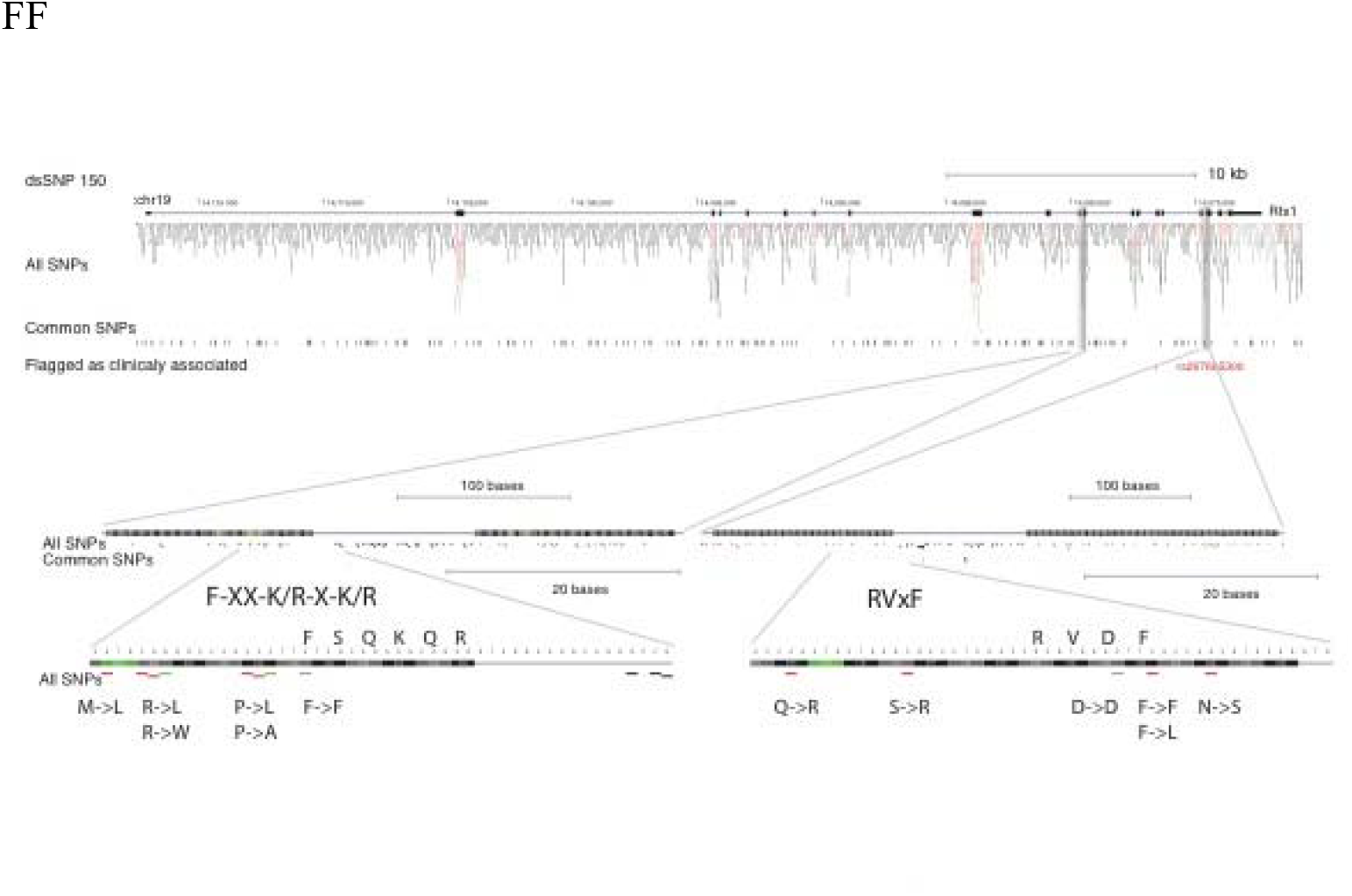
Browser image from the UCSC genome browser showing SNPs in the *RFX1* locus taken from the dbSNP 150 database. Common SNPs are SNPs found in at least 1% of samples. Red indicate non-synonymous change while green denote synonymous change in the amino acid sequence. The specific change for SNPs overlapping and adjacent to the PP1c binding motifs is indicated at the bottom.

### Supplementary Table 1

The SwissProt database was scanned for proteins containing the RVXF motif using the Expasy prosite search tool [2] (https://prosite.expasy.org/scanprosite/) using the pattern [RK]-[VI]-X-[FW] with the ineage/species filter = [Homo sapiens] (SuppData2.txt) and the description filter = [transcription] (SuppData3.txt).

### Supplementary Table 2

Enriched KEGG pathway for the transcription related proteins identified be prosite search. Analysis was preformed using the Strings website [3].

